# Treatment with saturating dose of conventional anti-CD28 monoclonal antibody well tolerated in pig acute myocardial infarction model

**DOI:** 10.64898/2026.01.15.699692

**Authors:** Anja Stadtmüller, Rebekka Grampp, Giulia Marianantoni, Florian Schnitter, Finja Becker, Könül Nuriyeva, Daniela Langenhorst, Leona Metka, Kira Günther, Lars Wiebusch, Kerstin H Mair, Nadine Gladow, Thomas Hermann, Jan Rohr, Ulrich Hofmann, Anna Frey, Niklas Beyersdorf

**Affiliations:** University Hospital Würzburg, Department of Internal Medicine I, Würzburg, Germany; University of Würzburg, Institute for Virology and Immunobiology, Würzburg, Germany; University Hospital Würzburg, Comprehensive Heart Failure Center, Würzburg, Germany; University of Veterinary Medicine Vienna, Immunology Unit, Centre of Pathobiology, Department of Biological Sciences and Pathobiology, Vienna, Austria; Albert-Ludwigs-University Freiburg, Institute of Immunodeficiency, Medical Center and Faculty of Medicine, Freiburg, Germany

**Keywords:** Myocardial infarction, pig, swine, heart, regulatory T cells, CD28, monoclonal antibody, non-superagonistic, inhibitor of ligand binding, immunotherapy

## Abstract

CD4^+^ Foxp3^+^ regulatory T cells (Treg) efficiently foster wound healing after myocardial infarction (MI). Therapeutically shifting the balance between CD4^+^ Foxp3^-^ conventional T cells (Tconv) and Treg towards Tregs enhanced survival in a mouse model of MI. Due to species-specific differences in cardiac wound healing and remodelling, it remains, however, unclear whether these findings can be translated into novel immunotherapies for human patients after MI. Therefore, we studied pigs whose cardiac wound healing after MI and the composition of the immune system are very close to humans. This includes the relevant complication of developing a cytokine release syndrome (CRS) after infusion of saturating amounts of a superagonistic anti-CD28 monoclonal antibody (mAb). To achieve the intended shift in the Treg/Tconv balance, we treated pigs three days after interventional MI induction with a non-superagonistic, i.e. conventional, anti-CD28 mAb, clone 3D11. Infusion of a saturating dose (1 mg/kg body weight) of mAb 3D11 was clinically well tolerated without signs of CRS induction or any other complications. Molecularly, mAb 3D11 infusion led to a downmodulation of CD28 expression on porcine T cells *in vivo* with the remaining CD28 molecules blocked from binding natural ligand proteins CD80 and CD86, as we show in this publication. Apart from modulating CD28 expression, treatment with mAb 3D11 did not induce any overt changes in the peripheral T cell compartment. However, after mAb 3D11 treatment, we observed Treg accumulation in the infarcted heart, particularly the border zone, on day 7 post-MI using immunofluorescence histology. Our findings thus suggest that even saturating doses of conventional anti-CD28 monoclonal antibodies could potentially be safely administered in patients to therapeutically shift the Treg/Tconv balance in the infarcted myocardium. This might be sufficient to enhance cardiac wound healing in patients short-term and prevent adverse remodelling long-term.

## Introduction

Myocardial infarction (MI) has a high prevalence and, when survived, often leads to chronic heart failure, which by itself comes with a high mortality (1). Whereas the therapeutic gold standard for MI is rapid restitution of coronary blood flow by percutaneous coronary intervention (1), unfortunately only 54% of patients undergo timely reperfusion (2). The remaining patients often suffer extensive transmural infarction, resulting in tissue necrosis and subsequent adverse myocardial remodelling (3). However, there are currently no immunotherapies available to modulate cardiac wound healing after MI and to prevent adverse remodelling despite considerable efforts in research and development (4). Beneficially modulating cardiac inflammation post-MI has the potential for a breakthrough in this field, as inflammation is one of the driving forces behind early healing as well as mid- and long-term remodelling processes (5). In rodent models of MI, we and others were able to establish that therapeutically shifting the balance of CD4^+^ Foxp3^-^ conventional (Tconv) and CD4^+^ Foxp3^+^ regulatory T cells (Treg) towards Treg fosters wound healing, reduces maladaptive cardiac remodelling and promotes long-term survival after MI (6, 7). This effect could be achieved either with a superagonistic anti-CD28 monoclonal antibody (CD28-SA) (7) or by injection of low-dose interleukin-2 (IL-2) (8, 9). Alternatively, blocking T cell costimulation with a conventional, i.e. non-superagonistic, anti-CD28 mAb blocking ligand binding (10) or with CTLA-4-Ig (Abatacept) (11) preferentially inhibited Tconv over Treg, also leading to a shift in the Treg/Tconv balance towards Treg and a better outcome after MI.

Compared to laboratory mice and rats, the human immune system faces many more challenges leading to ample formation of memory cells in the T and B cell compartments and a state of ‘training’ in cells of innate immunity (12). These differences pose significant challenges for the translation of immunotherapeutic approaches from rodents to humans. In case of MI, the sheer size of the human heart is another important difference to mice and rats, leading to markedly slower wound healing (13). Therefore, we assessed whether Treg activation after MI was feasible in conventionally housed pigs, i.e. in a human-like animal model. In a first study we used the superagonistic anti-porcine CD28 mAb 4D12 at a dose of 10 μg/kg body weight (bw) (14), i.e. well below the 100 μg/kg bw, which induces a cytokine release syndrome in humans (15) and pigs (16).

In this study, we now infused the conventional anti-pig CD28 mAb 3D11 (16) into pigs after MI at a saturating dose of 1 mg/kg bw. In-depth cardiological and immunological assessment of these animals until day 7 post MI revealed, on the one hand, no negative effects of mAb 3D11 infusion on cardiac function, while, on the other hand, treatment with mAb 3D11 led to an accumulation of Treg in the infarcted heart as determined by immunofluorescence histology.

## Material and Methods

### Animals and ethics statement

Pigs (German Landrace) were bred on Gerd Heinrichs’ farm (Heinsberg, Germany) and maintained in the animal facility of the Comprehensive Heart Failure Center Würzburg at the University Hospital of Würzburg. All animal experiments were conducted in accordance with the Animal Protection Law (Directive of the European Parliament and of the Council of 22 September 2010 (2010/63/EU)) and had been reviewed and approved by the review board of the District Government of Lower Franconia and the University of Würzburg (approval reference number 55.2.2.-2532-2-1402). The experiments were carried out in accordance with the ethical principles of the Declaration of Helsinki (1964, and later amendments) and the guidelines of the Federation of European Laboratory Animal Science Associations (FELASA). Surgical procedures were performed under general anaesthesia, and appropriate analgesia was provided to alleviate postoperative pain. Additionally, all efforts were made to minimise animal suffering and to reduce the number of animals used. C57BL/6J mice were bred and housed in the animal facility of the Institute for Virology and Immunobiology.

### Pig MI model

Pigs (aged 10 to 20 weeks with an average bw of 43 ± 7 kg) underwent MI induction under total intravenous anaesthesia with propofol (5 mg/kg bw/h i.v.) and fentanyl (12-24 µg/kg bw/h i.v.) according to the established protocol in our group (17). In brief, pigs were sedated with azaperone (8 mg/kg bw i.m.), including atropine (0.5 mg), anaesthetised with ketamine (15 mg/kg bw i.m.), intubated after a propofol bolus (2 mg/kg bw i.v.) and put on a mechanical ventilation for the duration of the procedure. After femoral artery puncture and subsequent coronary catheterisation, the pigs underwent temporary balloon occlusion of the left anterior descending artery (LAD) for 90 minutes (Supplemental Figure S1). Three pigs died during the LAD occlusion due to electrotherapy-resistant ventricular fibrillation despite continuous application of amiodarone (2 mg/kg bw/h i.v.) for arrhythmia prophylaxis. For additional pain prevention post-MI, the pigs received meloxicam (0.4 mg/kg bw i.m.) before completion of the anaesthesia at the end of the procedure and on the following day.

### Echocardiography

To analyse cardiac function, transthoracic echocardiography was performed on day 0 before MI (baseline), as well as on day 3 and on day 7 before termination of the experiment with an ultrasound system (Vivid E9; GE Healthcare, Chicago, IL, USA) under anaesthesia. Echocardiography was first performed from the right subxiphoid view (5-chamber), followed by left parasternal long- and short-axis views first in left and then in right lateral position. Parameters to assess cardiac function included stroke volume (LVOT diameter and curve), systolic and diastolic function (left ventricular ejection fraction using the Teichholz method in M-mode, E/A, E/É), and wall motion. Some recordings were acquired during short pauses of mechanical ventilation under mechanical ventilation to improve image quality, which was not feasible in spontaneously breathing animals. The evaluation of parameters was performed in a post-processing manner by a blinded doctoral researcher. All parameters were measured three times for each animal and the arithmetic mean was used for comparative analysis between the groups.

### Characterisation of mAb 3D11 *in vitro*. Binding of porcine CD80 to porcine CD28

HEK-293T cells were transfected with a plasmid encoding for a fusion protein of the extracellular domain of porcine CD80 and the Fc part of human IgG_1_ (pCD80-hIg). Prior to transfection, the plasmid was sequenced. Supernatant of the transfected cells was incubated with A20J cells expressing porcine CD28 or wild-type A20J cells as a control. Bound pCD80-hIg was visualised using anti-human Ig-Alexa647 (Jackson ImmunoResearch Europe Ltd., Ely, United Kingdom). A20J were preincubated with mAb 3D11 before pCD80-hIg was added to assess blocking of ligand binding by mAb 3D11.

### Trogocytosis (transendocytosis) of mouse CD80 by porcine CD4^+^ T cells

Porcine splenocytes were seed on top of mouse embryonic fibroblasts (MEFs) expressing mouse CD80 tagged with the mScarlet fluorophore to study trogocytosis following published protocols (18, 19). Wild-type MEFs were used as a control. After four hours, porcine splenocytes were resuspended and stained for CD4 and CD8 expression and the median fluorescence intensity (MFI) of mScarlet was determined on CD4^+^ CD8^-^ splenocytes by flow cytometry.

### Trogocytosis (transendocytosis) of mouse CD86 by porcine CD4^+^ T cells

Mouse and porcine splenocytes were co-cultured overnight and the stained for porcine CD4 (PE, BD and Co., Franklin Lakes, NJ, USA.) and CD8 (FITC, BD) as well as mouse CD86 expression (biotin - Streptavidin-PE-Cy5, both BD). The MFI of the bound anti-mouse CD86 was determined on CD4^+^ CD8^-^ porcine splenocytes by flow cytometry.

### Antibody infusion

On day 3 after MI, pigs were again sedated and intubated as described above, but were now maintained on inhalation anaesthesia with isoflurane (1–2 vol%) to preserve spontaneous breathing and facilitate recovery. In advance, the animals had independently and randomly been assigned to either the treatment group receiving mAb 3D11, (InVivo BioTech Services GmbH, Henningsdorf, Germany; <1 EU/mg Endotoxin level), or to the control group receiving isotype control antibody at the same dosage of 1 mg/kg bw (MOPC-21; Bio X Cell, Inc., Lebanon, NH, USA; <1 EU/mg Endotoxin level). The agent was provided in a blinded manner in 10 ml 0.9% sterile saline solution and then further diluted in 500 ml 0.9% sterile saline. The total volume of 510 ml was administered i.v. within 30 minutes under continuous cardiopulmonary monitoring. 11 out of 12 pigs survived the antibody infusion and exhibited no signs or symptoms of an allergic reaction or even a CRS. One animal died during the mAb 3D11 infusion, most likely due to acute volume overload leading to a clinical presentation of a cardiogenic shock, however, an acute reaction to mAb 3D11 cannot be completely ruled out. The researchers who performed the MI, and the echo, applied the infusion and conducted the subsequent processing of the harvested organs were blinded to the antibody infused to obtain unbiased results. Unblinding for infused therapy occurred only after all reported individual measurements had been evaluated and the analysis of the whole data set had started.

### Serial blood sampling and preparation of PBMC

Blood samples were taken on day 0, i.e. before MI induction, as well as before the procedures on days 3 and 7. Serum samples were centrifuged, the supernatant was collected and preserved in aliquots at −80 °C. Furthermore, heparinised blood samples were taken for whole blood analysis and for peripheral blood mononuclear cell (PBMC) isolation via the Ficoll separation method (Histopaque-1077; Sigma-Aldrich, St. Louis, MO, USA). The isolated PBMC were resuspended in cell culture medium RPMI-1640 containing 10% fetal calf serum (FCS; Gibco™, Thermo Fisher Scientific, Waltham, MA, USA), 2-Mercaptoethanol, sodium pyruvate, MEM solution, 5% Glutamine, Streptomycin, Penicillin and counted. 1×10^6^ cells (2×10^6^ on day 7) were further stained for flow cytometry and the remaining cells were preserved in cell cryomedium (RPMI-1640, FCS, Dimethylsulfoxid – DMSO) at −80 °C.

### Organ preparation on day 7 post MI

On day 7 after MI anaesthesia was induced as described above. Blood samples were taken from the animals and the final echocardiography was completed. After bolus administration of heparin i.v. (300 IU/kg bw) pigs were euthanised with pentobarbital i.v. (dosage according to manufactureŕs instructions) for immediate organ extraction. Parts of the spleen, thymus and mediastinal lymph nodes were removed and submerged in cold RPMI-1640 medium. From there, the samples were cut into smaller pieces, weighed and grounded through a 70 μm cell strainer into cold Hankś Balanced Salt Solution without Ca^2+^ and Mg^2+^, with phenol red and with 0.1% bovine serum albumin (HBSS -/- /BSA). With the Ficoll separation method, splenocytes and thymocytes were isolated. After a final centrifugation, all the cell suspensions were resuspended in a cell culture medium and stored overnight at 4 °C. On the next day, the cells were counted and either further stained for flow cytometry or preserved in cell cryomedium.

After removal, the hearts were washed with ice-cold 0.9% saline. In addition, the coronaries on both sides were manually flushed to remove any blood residues. The organ was weighed and photographed. The left ventricle was then cut into evenly thick transverse slices (8-10 mm), which were weighed and photographed individually. Samples of 600 mg tissue from the different regions, i.e. infarct core (IC), border zone (BZ), and remote myocardium (RM), were taken from infarcted heart slices. At least two different sites were sampled per region - the respective regions had been selected purely macroscopically (Supplemental Figure S1). The specimens were next processed for immunofluorescence staining and flow cytometry.

To isolate leukocytes, the heart tissue was finely chopped and digested at 37 °C for 30 minutes with DNAse I (60 units/ml Roche Diagnostics GmbH; Mannheim, Germany) and collagenase type 2 (600 units/ml Worthington Biochemical, Lakewood, NJ, USA) in Hanks’ balanced salt solution (HBSS +/+, with Ca^2+^/Mg^2+^ without phenol red). The samples were then first passed through 70 µm and then 40 µm cell strainers in cold HBSS -/- /BSA. This was followed by 8 minutes of red blood cell lysis with an RBC-lysis buffer (10x, BioLegend, San Diego, CA, USA). To identify viable cells, live-dead staining with Zombie Aqua 1:200 (fixable viability kit (DMSO); BioLegend, San Diego, CA, USA) was carried out in DPBS−/− /EDTA (Dulbecco’s Balanced Salt Solution without Ca^2+^/Mg^2+^, containing 2 mM ethylene diamine tetra-acetic acid) at room temperature, in the dark, for 15 minutes. After washing the samples with flow cytometry buffer (DPBS−/− /EDTA with 2 % FCS), the cells were blocked with pig serum 10 % (own production; in DPBS−/− /EDTA) for 15 minutes at 4 °C. The next step was the anti-CD45-FITC antibody staining (1:50, Mouse anti Pig CD45 FITC; Bio-Rad; clone: K252.1E4) for 30 minutes at 4 C. To ensure a sufficient yield of leukocytes from all three areas of the heart from which we took samples (IC, BZ, RM), magnetic activated cell sorting (MACS) was performed. For that, after washing the cells with flow cytometry buffer, they were incubated for 15 minutes at 4 °C with anti-FITC MicroBeads (1:20, Miltenyi Biotec, Bergisch Gladbach, Germany). Another washing step followed and CD45^+^ cells were positively isolated using LS columns (Miltenyi Biotec, Bergisch Gladbach, Germany) according to the manufacturer’s instructions.

### Flow cytometry. Whole blood samples

50μl of whole blood were transferred to TruCOUNT tubes containing TruCount Beads (BD Biosciences, Franklin Lakes, NJ, USA) and processed following the manufacturer’s instructions - including calculation of absolute cell numbers. Antibodies used for staining are listed in Supplemental Table 1. Whole blood samples were analysed on a FACS Calibur (BD FACSCalibur^TM^ Flow Cytometer, BD Biosciences). Isolated PBMC were stained with the antibody panel summarised in Supplemental Table 2.

### Cardiac leukocytes, mediastinal lymph nodes, spleen and PBMCs

Antibodies used for staining cardiac lymphocytes, cells from mediastinal lymph nodes, spleen and PBMCs are listed in Supplemental Tables 2 - 3. The fixation and permeabilisation kit (eBioscience Foxp3/Transcription Factor Staining Buffer Set - Thermo Fisher Scientifc, Carlsbad, CA, USA) was used to intracellularly stain Foxp3 und Ki-67 (see Supplemental Table 3, staining 5-7). Supplemental Figure S2 shows an example of our gating strategy.

Data analysis was conducted using FlowJo v10.10.0 (FlowJo, Ashland, OR, USA) following measurements obtained with the Attune NxT Flow Cytometer (Thermo Fisher Scientific, Waltham, MA, USA) and Calibur.

### Immunofluorescence histology of cardiac tissue

For immunofluorescence staining samples of cardiac tissue from different regions (IC, BZ, RM) were embedded in cryomedium (Tissue-Tek; Sakura Finetek Europe, Alphen aan den Rijn, The Netherlands) on dry ice and preserved at −80 °C. A few days before staining, the frozen tissue was cut into 7 µm thick slices with a cryostat microtome (CM1850; Leica Biosystems, Wetzlar, Germany), applied to microscope slides and retained at −20 C. For subsequent staining, the cryosections were thawed for 30 minutes at room temperature and fixed in 4% PFA (paraformaldehyde). Then, the slides were washed in tris-buffered saline (TBS) (3 x for 5 minutes) and permeabilized with 0.02% Triton (Triton X-100) for 10 minutes. Washing steps were repeated, followed by the application of the blocking medium (60 minutes – IHC/ICC Blocking Buffer High Protein, Invitrogen) in a humid chamber at room temperature. Afterwards, anti-CD4 PE (Mouse anti Pig CD4 Alpha RPE; Bio-Rad; clone MIL17) was applied (1:50 diluted in blocking medium) and the samples were stored at 4 °C overnight (continuously in the humid camber). On the following day, the washing routine was changed to initiate the buffer switch [2 x 5 minutes in TBS and 1 x 5 minutes in phosphate-buffered saline (PBS)]. Next, the permeabilization with 0.5% dodecyltrimethylammonium-chloride (DTAC) for 20 minutes took place and was followed by staining with anti-Foxp3 APC (monoclonal antibody; eBioscience™; Invitrogen; clone FJK-16s) 1:200 diluted in blocking medium at 4 °C overnight in the humid chamber. Lastly, the slides were washed with PBS (3 x 5 minutes) and the cell nuclei were stained with DAPI (4′,6-diamidino-2-phenylindole – 1:5000) for 10 minutes at room temperature in the humid chamber. The washing steps were repeated and the samples were mounted with ImmuMount (Epredia Netherlands B.V., DA Breda, Netherlands). For analysis, the DMi8 fluorescence microscope (Leica Microsystems, Wetzlar, Germany) was used. The sections were examined manually using ImageJ. At least 100 CD4+ cells should be counted, but no more than 6 visual fields (715µm x 530µm) were considered. For the BZ definition, both the IC and the RM had to be visible in the selected field. CD4^+^ cells and double-positive CD4^+^ Foxp3^+^ cells were assessed separately. The cell counts were then put into relation to the number of counted visual fields.

### Statistical data analysis

Statistical analysis and data visualisation were performed using GraphPad Prism v10.6.0 (GraphPad Software, San Diego, CA, USA). Values for each animal are shown individually with horizontal bars indicating means per group. Where only means and standard deviations are shown, the number of samples are specified in the figure legends. The statistical tests applied are also mentioned in the figure legends. A p value < 0.05 was considered statistically significant.

## Results

### Monoclonal antibody 3D11 blocks ligand binding to porcine CD28

The anti-porcine CD28 mAb 3D11 is a conventional anti-CD28 mAb that recognises an epitope distinct from that bound by the superagonistic anti-CD28 mAb 4D12 (16). To test whether mAb 3D11 also blocks ligand binding to porcine CD28, we incubated A20J cells (mouse B cell line) recombinantly expressing porcine CD28 with a fusion protein of the extracellular domain of pigCD80 and the Fc part of human IgG_1_ (pigCD80-hIg) in the presence or absence of mAb 3D11. Binding of pigCD80-hIg to porcine CD28 was diminished in the presence of mAb 3D11 (Figure 1A, B).

**Figure 1.**
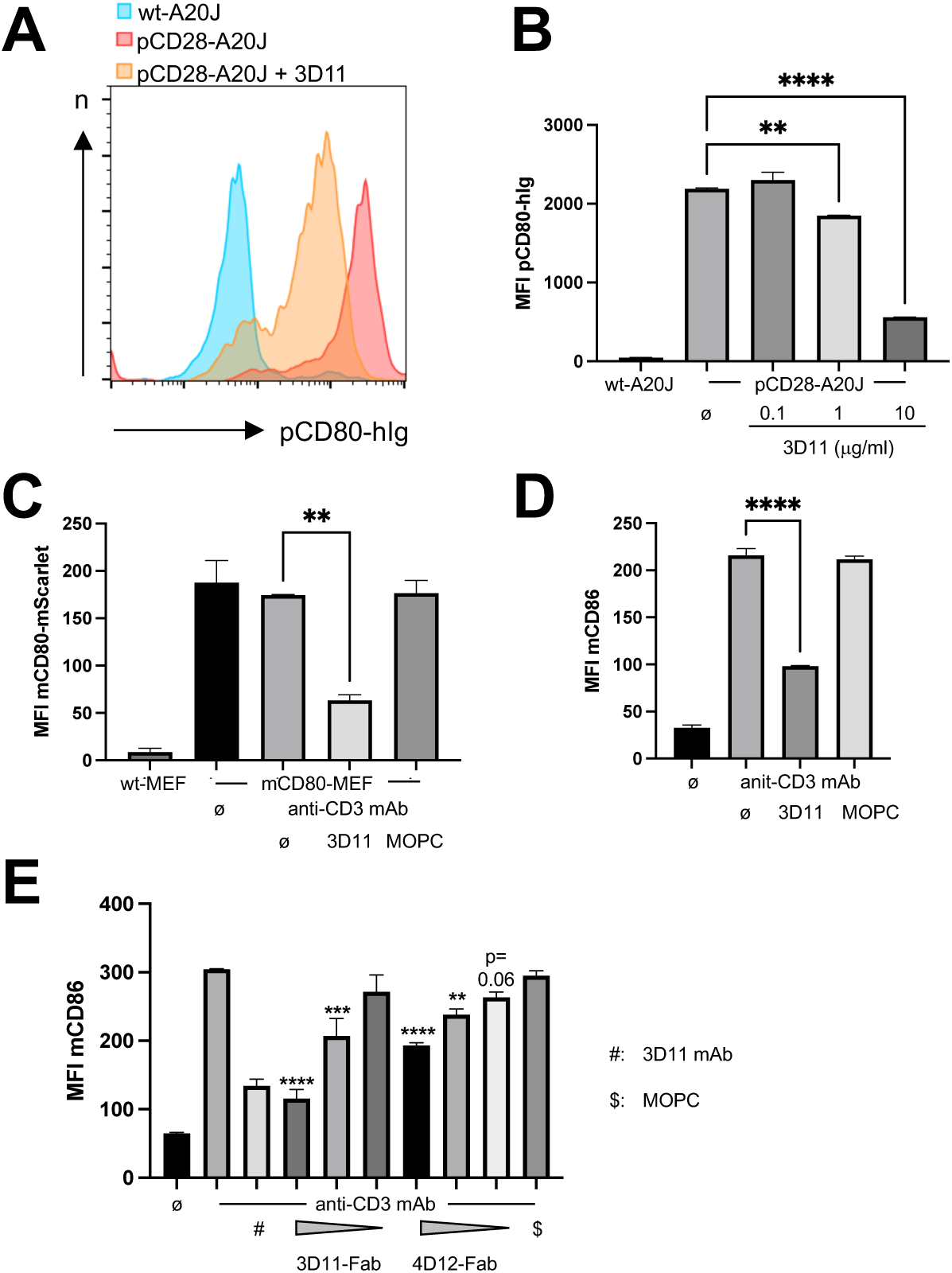
Anti-pig CD28 mAb 3D11 blocks ligand binding. **A**) Representative histograms showing pCD80-hIg binding to wt or pCD28-transgenic A20J cells in the absence or presence of 3D11 mAb. **B**) Summary graph of MFI of pCD80-hIg under the condi-tions shown. **C**) MFI of mouse CD80-mScarlet on porcine CD4^+^ T cells after trogocytosis from transgenic MEFs (4 h incubation). **D**) MFI of mouse CD86 on porcine CD4^+^ T cells after trogocytosis from mouse splenocytes (incubation overnight). **E**) Set-up as for D). C)-E): Anti-CD3 mAb: 1 μg/ml mAb PPT-3. 3D11 and MOPC-21 mAb: 10 μg/ml. Concentrations Fab fragments: 20, 5, 1.25 μg/ml. B)-E): Means + SD of duplicates. Data representative of at least two experiments with similar results. One-way ANOVA followed by Tukey’s post-hoc test (Holm-Sidak in E). ** p < 0.01, *** p < 0.001, **** p < 0.0001.

Recently, it has been described that trogocytosis (transendocytosis) of its ligands is an integral part of the biological function not only of CTLA-4, but also of CD28 (20, 21). Therefore, we next assessed the impact of mAb 3D11 on trogocytosis of CD80 and CD86 by porcine CD4^+^ T cells. For mouse embryonic fibroblasts (MEFs) expressing mouse CD80 tagged with mScarlet (mCD80-mScarlet), we noted that porcine CD4^+^ T cells acquired mCD80-mScarlet without CD3 ligation (Figure 1C). This ‘spontaneous’ trogocytosis of mCD80-mScarlet was inhibited by mAb 3D11, but not by the control mAb MOPC-21. In a second experimental system, we used mouse splenocytes as a source for the CD28 ligand CD86. Here, ligation of CD3 on porcine CD4^+^ T cells induced trogocytosis of mouse CD86 (Figure 1D), which was again inhibited in the presence of mAb 3D11. As binding of mAb 3D11 to porcine CD28 by itself induces T cell costimulation, we generated Fab fragments of mAb 3D11 to study inhibition of costimulation by blockade of ligand binding to CD28.

For comparison, we also generated Fab fragments of the superagonistic anti-CD28 mAb 4D12. As expected, 3D11-Fab was much more potent than 4D12-Fab at inhibiting trogocytosis of CD86 by porcine CD4^+^ T cells (Figure 1E). Moreover, 3D11-Fab and, again, much less 4D12-Fab, inhibited the proliferation of CD4^+^ porcine splenocytes stimulated with Concanavalin A (Con A; Figure 2). This inhibition could be overcome by adding the anti-CD28 mAb 4D12 together with 3D11-Fab (Figure 2A and 2B ‘#’) confirming that the diminshed proliferation seen in the presence of 3D11-Fab was truly due to the inhibition of CD28 costimulation.

**Figure 2.**
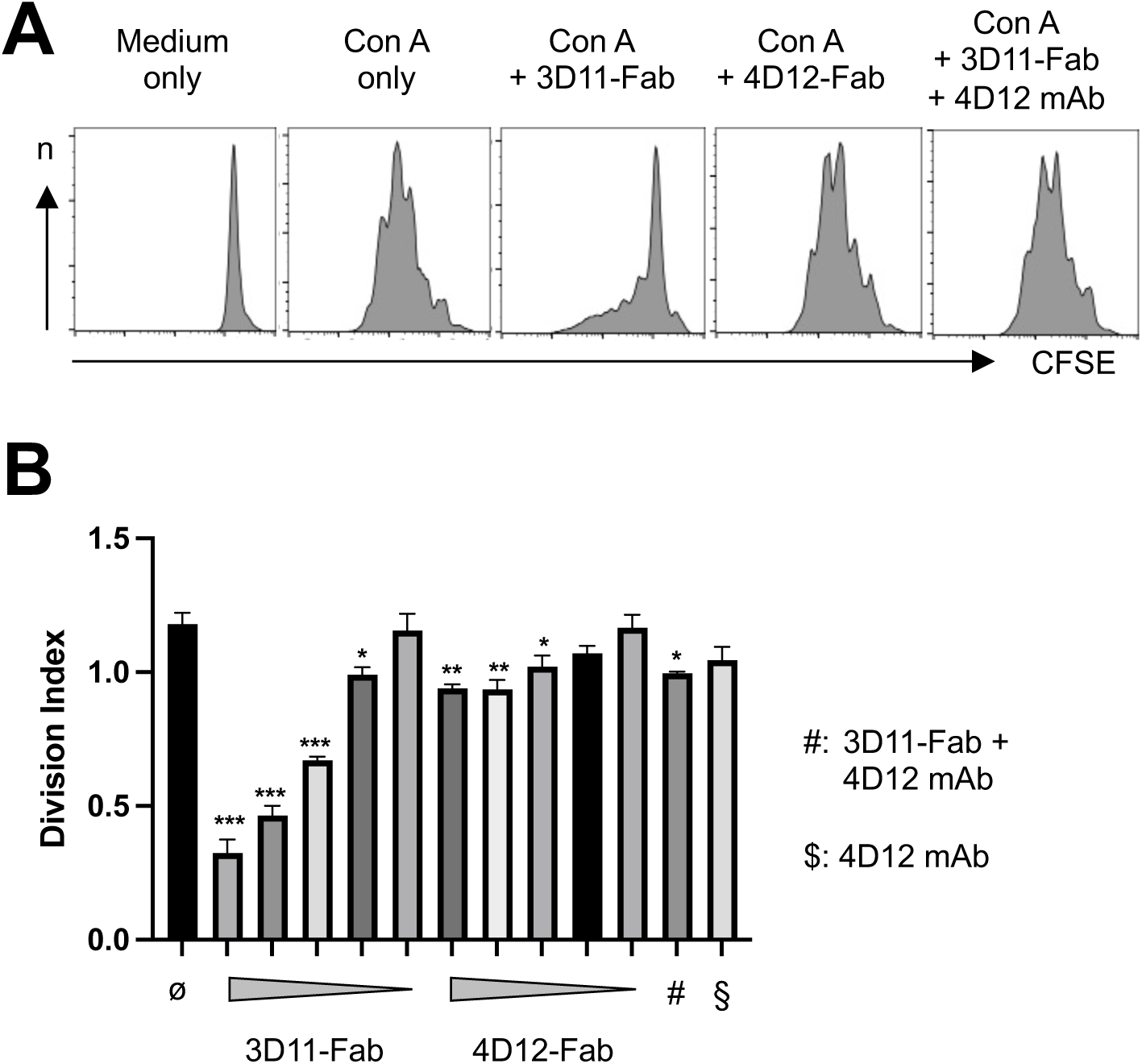
Fab fragments of 3D11 mAb inhibit T cell costimulation. CFSE-labelled porcine splenocytes were stimulated with Con A for four days in the presence or absence of Fab fragments of anti-pCD28 mAb 3D11 or 4D12 (20 μg/ml each). **A**) Representative CFSE dilution profiles of gated CD4^+^ cells. **B**) The FlowJo^TM^ proliferation tool was used to determine division indices. Means + SD of duplicate cultures. Concentrations of Fab fragments: 20, 10, 5, 2.5 and 1.25 μg/ml. Concentration of 4D12 mAb: 10 μg/ml. #: 20 μg/ ml 3D11-Fab plus 10 μg/ ml 4D12 mAb. One-way ANOVA followed by Holm-Sidak test. * p <0.05, ** p < 0.01, *** p < 0.001: Comparison with cells cultured with Con A only. The experiment was repeated three times with similar result.

Blockade of ligand binding to porcine CD28 together with the inhibitory activity of Fab fragments of 3D11 on T cell proliferation, identify mAb 3D11 as a ligand-binding blocking anti-CD28 mAb.

### Infusion of mAb 3D11 did not affect the deterioration of cardiac function within the first week after MI

Both in the pig MI model (22) as well as in humans (1) with similarly large infarcts, cardiac function continues to deteriorate in the days following MI. To determine whether mAb 3D11 infusion (1 mg/kg bw) has a short-term impact on cardiac function, we conducted serial echocardiography on pigs before and after MI induction. For these analyses, we included data from a total of eleven pigs (seven treated with mAb 3D11 and four treated with the control mAb MOPC - also 1 mg/ kg bw) that survived the whole observation period. In echocardiography, reductions in the left ventricular ejection fraction (LV-EF, Figure 3D) and the fractional shortening (LV-FS, Figure 3E) clearly indicated the expected deterioration in cardiac function during the first week after MI. In addition, abnormal motion of both the septal and the anterior walls of the left ventricle confirmed successful induction of MI (data not shown) (14). However, there were no statistically significant differences between conventional anti-CD28 mAb-treated pigs and pigs that had received the control mAb MOPC in any of the parameters analysed (Figure 3). The echocardiographic data, thus, showed that infusion of 1 mg/ kg bw of the conventional anti-CD28 mAb 3D11 had no negative side-effects on cardiac function in any of the infarcted pigs. The short observation period and the early timepoints after MI analysed might have precluded observing positive effects of mAb 3D11 infusion on cardiac remodelling as might be seen long-term and as previously observed in mice (7).

**Figure 3.**
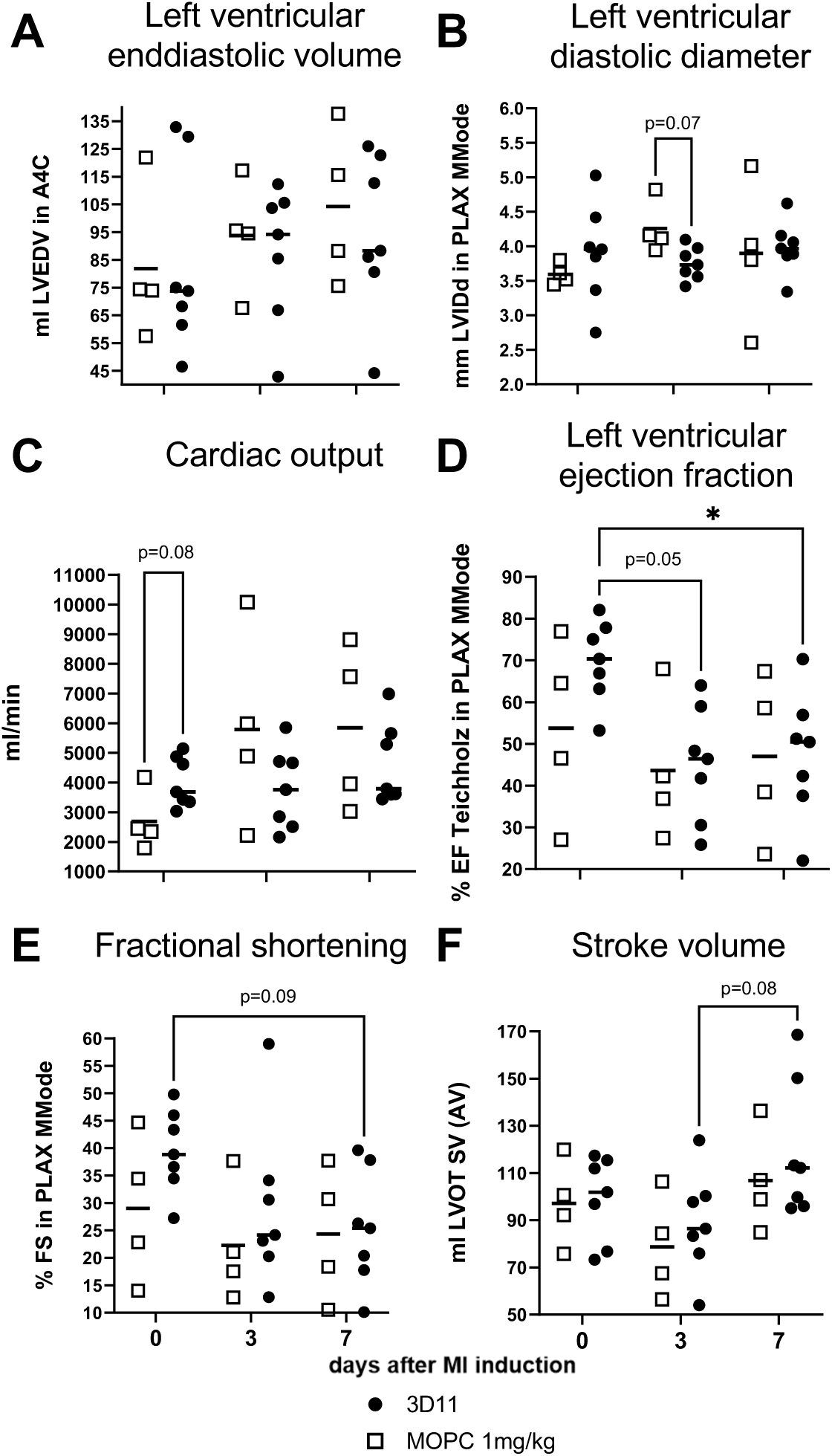
Similar cardiac function as determined by echocardiography in pigs on day 7 after MI, irrespective of antibody treatment. mAb 3D11 was infused on day 3 after MI induction. **A**) Left ventricular enddiastolic volume (LVEDV) in the apical 4-chamber view (A4C) was similar on the day of MI induction as well as on days 3 and 7 after MI without differences between both groups. **B**) Left ventricular internal diastolic diameter (LVIDd) in parasternal long-axis view (PLAX) is stable over time after MI without differences between both groups. **C**) No differences in cardiac output over time and between both groups. **D**) Left ventricular ejection fraction (EF) by the Teichholz method in PLAX and **E**) fractional shortening (FS) in PLAX decreased over time due to MI induction without differences between both groups. **F**) Stroke volume (SV) by LVOT diameter and LVOT curve is stable over time after MI without differences between both groups. Repeated measures two-way ANOVA followed by Tukey’s post-hoc test. numbers. * p < 0.05.

### Downmodulation of CD28 expression on T cells in peripheral blood and secondary lymphoid organs after infusion of saturating amounts of mAb 3D11

For the identification and characterisation of mAb 3D11 as a conventional anti-CD28 mAb blocking ligand binding, *in vitro* studies were sufficient (16) (Figure 1, 2). To study the *in vivo* activity of mAb 3D11 in a therapeutic setting, we infused mAb 3D11 or control mAb MOPC-21 at 1 mg/ kg body weight (bw) into pigs three days after MI induction. On day seven after MI induction, mAb 3D11 was still readily detectable on CD4^+^ T cells from peripheral blood (Figure 4A), spleen and mediastinal lymph nodes (Figure 5A). Moreover, CD28 expression was decreased on CD4^+^ T cells after mAb 3D11 infusion (Figure 4B, 5B). Comparison of the signals obtained from *in vivo*–infused mAb 3D11 (Figure 4A, 5A) and from staining for CD28 expression using unconjugated 3D11 as primary antibody (Figure 4B, 5B) indicated complete saturation of CD28 by mAb 3D11. Neither MI induction nor antibody infusion had an impact on the absolute numbers of total T cells or of CD4^+^ and CD8^+^ T cells in peripheral blood (Figure 4C - E). The same was true for the proportion of CD4^+^ T cells among leukocytes of spleen, mediastinal lymph nodes (Figure 5C) and blood (Figure 4F) and the proportion of Treg among CD4^+^ T cells (Figure 4G, 5D). The proportion of proliferating Ki-67^+^ cells among Treg and Tconv of peripheral blood (Figure 4H, I), spleen and mediastinal lymph nodes (Figure 5E, F) were also similar in 3D11- and MOPC-treated pigs. Together, the main effect of infusion of mAb 3D11 at 1 mg/kg bw was to reduce CD28 expression on T cells about fivefold and to achieve saturated binding of mAb 3D11 to the remaining CD28 molecules.

**Figure 4.**
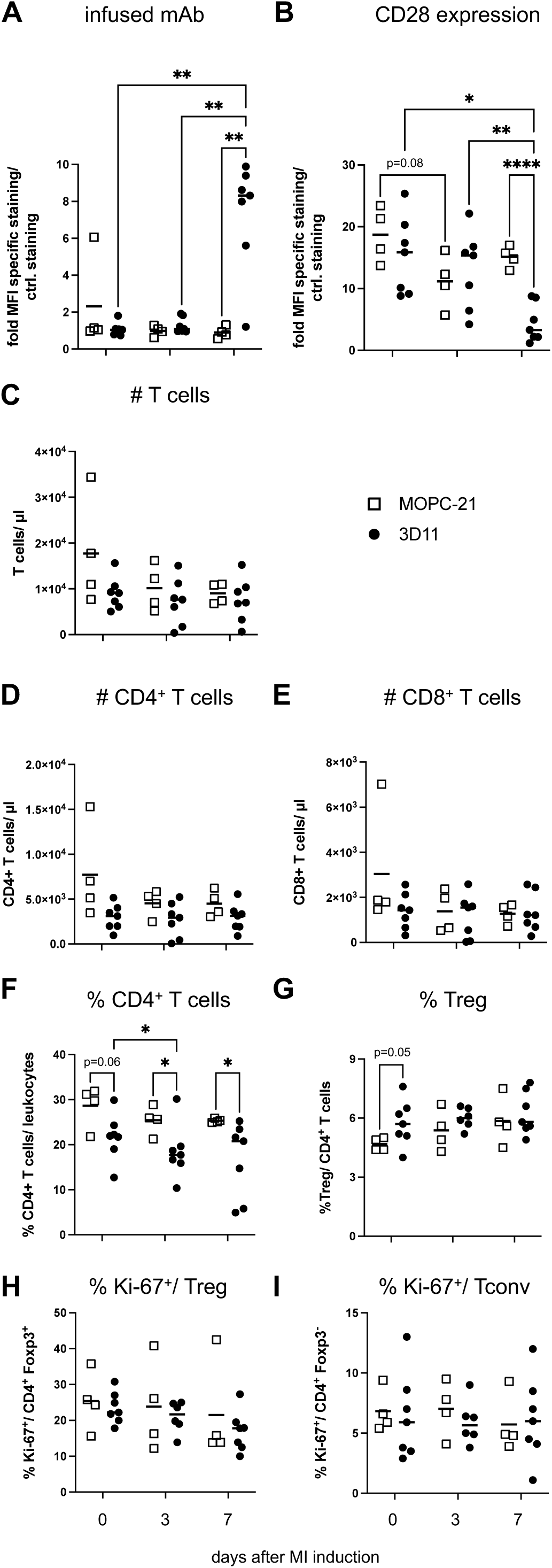
Tracking of infused mAb 3D11 in vivo and longitudinal analysis of T cell subsets in peripheral blood. mAb 3D11 was infused on day 3 after MI induction. **A**) Detection of mAb 3D11 on CD4^+^ T cells of peripheral blood on the day of MI induction as well as on days 3 and 7 post MI induction. **B**) CD28 expression by peripheral-blood CD4^+^ T cells. **C**) Absolute number of total T cells, **D**) CD4^+^ T cells and **E**) CD8^+^ T cells per μl of blood. **F**) Proportion of CD4^+^ T cells among peripheral-blood leukocytes. **G**) Proportion of CD25^+^ Foxp3^+^ Treg among CD4^+^ T cells. **H**) Proportion of Ki-67^+^ cells among Treg and **I**) Tconv. Repeated measures two-way ANOVA followed by Tukey’s post-hoc test. 0.05 ≤ p < 0.1 are shown as numbers. * p < 0.05, ** p < 0.01, *** p < 0.001.

**Figure 5.**
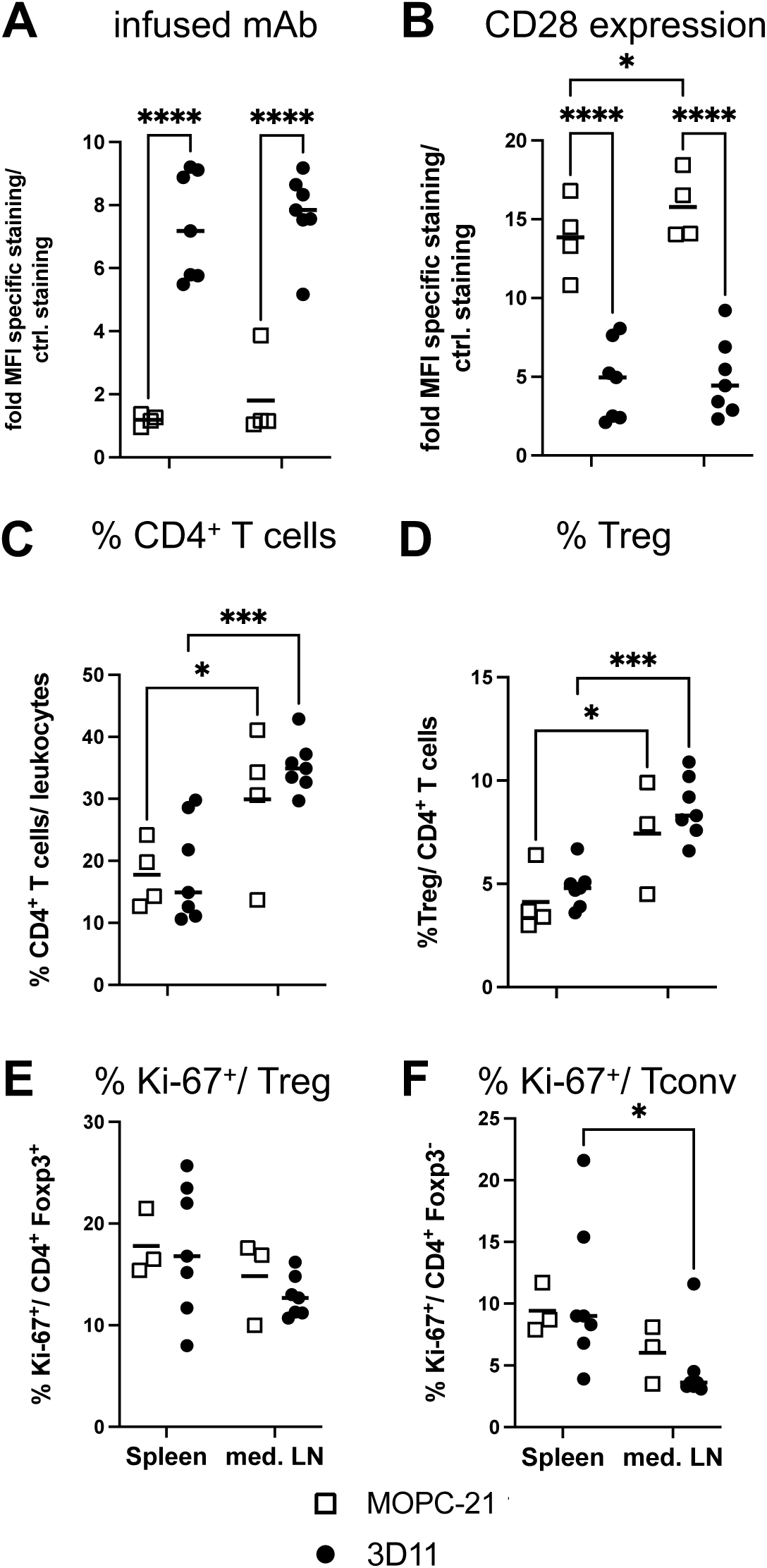
mAb 3D11 infusion decreased CD28 expression on CD4^+^ T cells of the spleen and mediastinal lymph nodes on day 7 after MI. **A**) Detection of mAb 3D11 on CD4^+^ T cells. **B**) CD28 expression by CD4^+^ T cells. **C**) Proportion of CD4^+^ T cells among leukocytes. **D**) Proportion of CD25^+^ Foxp3^+^ Treg among CD4^+^ T cells. **E**) Proportion of Ki-67^+^ cells among Treg and **F**) Tconv. Repeated measures two-way ANOVA followed by Tukey’s post-hoc test. * p < 0.05, *** p < 0.001, **** p < 0.0001.

### Infusion of mAb 3D11 three days after MI induction increased proportion of Treg among heart-infiltrating CD4^+^ T cells on day seven after MI induction

To understand how treatment with mAb 3D11 affected heart-infiltrating CD4^+^ T cells, we isolated CD45^+^ leukocytes from the infarcted myocardium to further analyse the CD4^+^ T cell compartment by flow cytometry. However, we were only able to successfully retrieve sufficient numbers of leukocytes for flow cytometry from four mAb 3D11- and three MOPC-21-treated animals. The data suggest that, as for peripheral blood, spleen and mediastinal lymph nodes, there were no changes in the proportion of CD4^+^ T cells among total leukocytes (Figure 6A), in the proportion of Treg among heart-infiltrating CD4^+^ T cells (Figure 6B) or in the proportion of Ki-67^+^ cells among cardiac Treg (Figure 6C) and Tconv (Figure 6D) after infusion of mAb 3D11. For immunofluorescence histology material from three MOPC-21- and all seven 3D11-treated pigs was available. Here, we observed higher numbers of Treg in the infarcted versus the remote myocardium, particularly the border zone, after mAb 3D11 infusion (Figure 6E), which was also the case for Tconv (Figure 6F). Autofluorescent signals detected in the green channel allowed us to separate the infarct core from the border zone and the remote myocardium (data not shown) (14). Moreover, for CD4, we observed a cell surface staining, while the Foxp3 signal overlaid with the DAPI signal, indicating nuclear localisation (data not shown) (14). Thus, we observed an accumulation of Treg in the infarcted myocardium after treatment with mAb 3D11.

**Figure 6.**
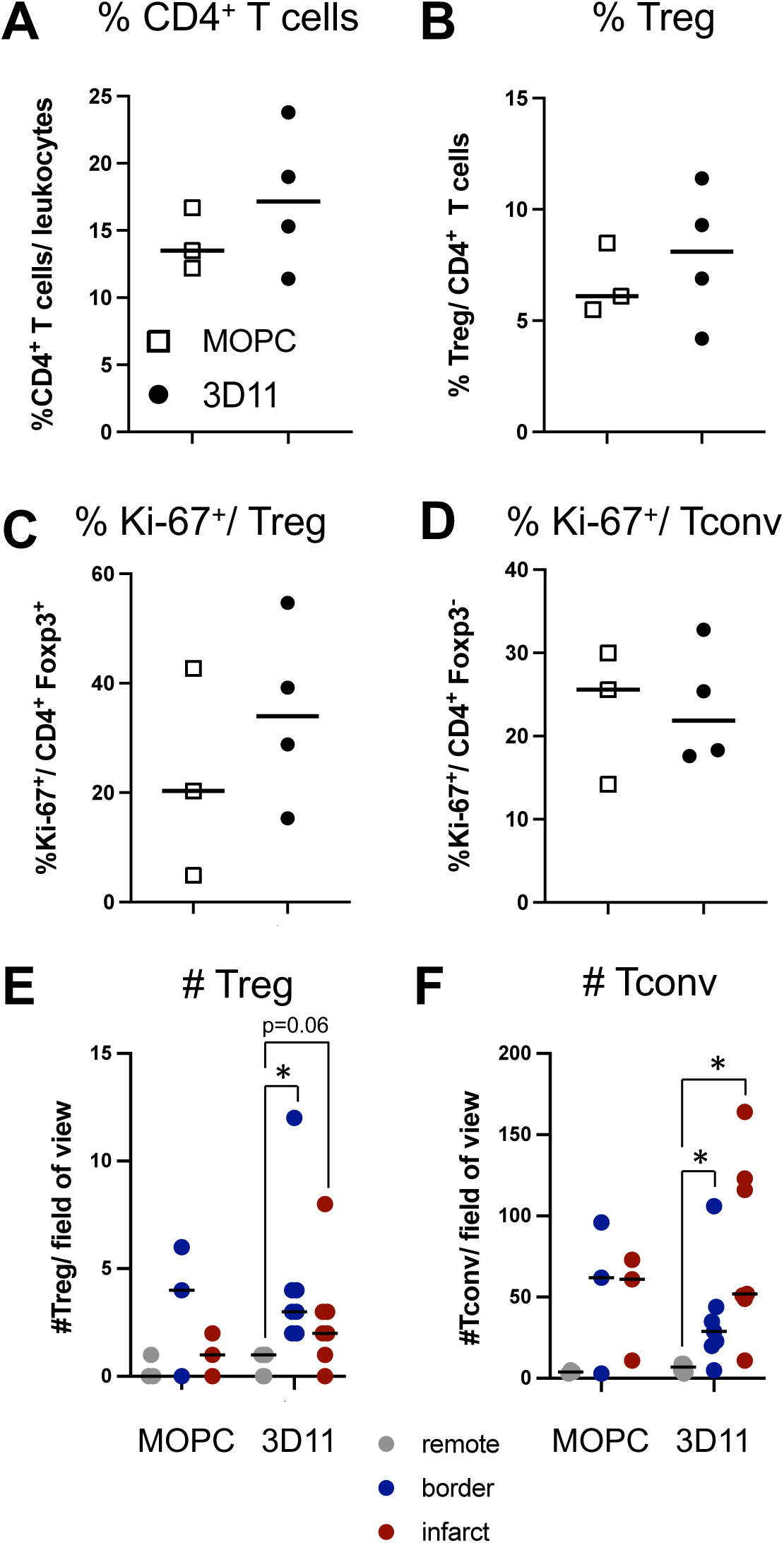
Elevated numbers of Treg in the border zone after mAb 3D11 infusion. **A**) Proportion of CD4^+^ T cells among cardiac leukocytes. **B**) Frequencies of Treg among cardiac CD4^+^ T cells. **C**) Proportion of Ki-67^+^ cells among cardiac Treg and **D**) Tconv. **E**) Tissue sections of infarcted hearts were stained for CD4 and Foxp3 expression. Treg and **F**) Tconv per field of view (715 um x 530 um). Up to six fields of view were analysed. Conventional anti-CD28 mAb 1 mg/ kg bw; MOPC-21: control mAb. Stats: Two-tailed paired Wilcoxon test; * p < 0.05.

## Discussion

In this work, we describe the short-term immunological and cardiological effects of infusion of a saturating dose of the conventional anti-CD28 mAb 3D11 in pigs recovering from myocardial infarction. Our data show that infusion of mAb 3D11 was well tolerated by the animals and that it did not negatively affect cardiac function during the first week after MI. Within the peripheral T cell compartment, our main observation was the downmodulation of CD28 expression after infusion of mAb 3D11 with no obvious impact on the Treg/Tconv balance in flow cytometry. In the infarcted heart, however, Treg accumulated in pigs treated with mAb 3D11 as shown by histological analysis. This work is, thus, the first to show that conventional anti-CD28 mAb can be administered safely even at a saturating dose with the aim to therapeutically manipulate cardiac Treg accumulation in a species prone to develop a CRS after high-dose CD28-SA infusion (16).

Generally, conventional anti-CD28 mAb bind epitopes outside of the so-called C’’D loop recognised by superagonistic anti-CD28 mAb (23). This means that conventional anti-CD28 mAb harbour clones which bind to or close to the binding site of CD80 and CD86 on the tip of the CD28 molecule and are, thus, capable of blocking ligand binding to CD28. Using pCD80-hIg binding for screening anti-porcine CD28 mAb generated by us, we observed that about half of the 25 clones analysed at least partially blocked pCD80-hIg binding to pCD28-expressing A20J cells with 3D11 being among the most efficient (data not shown).

After *in vivo* infusion, mAb 3D11 not only bound to CD28, but also led to its downmodulation. Reduced CD28 expression is most likely a consequence of agonism of mAb 3D11 on CD28 leading to CD28 internalisation via clathrin-coated pits (24). About two-thirds of CD28 expression were lost from the T cell surface after mAb 3D11 infusion (Figure 3, 4). This might reflect a similarly graded accessibility of CD28 molecules towards anti-CD28 mAb binding in pigs as in mice where CD28 is pre-arranged in clusters on the T cell surface (25). Alternatively, re-expression of CD28 molecules on the T cell surface by the day of analysis, i.e. four days after mAb 3D11 treatment, might explain retention of CD28 expression at reduced levels.

CD28 molecules expressed at the cell surface and bound by mAb 3D11 can neither be engaged by CD80 or CD86 to mediate costimulation of the T cells nor are they able to maintain the availability of CD28 ligands through trogocytosis (Figure 1) and re-expression at the cell surface (20, 21). This means that CD80 and CD86 molecules remain exposed to CTLA-4-mediated trogocytosis and degradation, potentially constituting an indirect form of immunoregulation mediated by mAb 3D11.

Retaining the biological activity of CTLA-4 is a major advantage of selectively blocking CD28 versus blocking both CD28 and CTLA-4 (26) as is the case for CTLA-4-Ig (Abatacept) which binds to CD86/ CD80-expressing antigen-presenting cells (27). In a Baboon model of memory T cell–driven skin inflammation, selective blockade of CD28 reduced symptoms, while infusion of CTLA-4-Ig had no effect (28). While, of course, in e.g. patients with rheumatoid arthritis the clinical value of treatment with CTLA-4-Ig is well documented (29), no data are currently available concerning the efficacy of CTLA-4-Ig in patients after MI.

Despite infusing a saturating amount of mAb 3D11, no changes to the absolute numbers of T cells in peripheral blood or their representation in secondary lymphoid organs (Figures 4, 5) indicate that the antibody neither induced complement nor antibody-dependent cellular cytotoxicity (CDC, ADCC). Thus, mAb 3D11, indeed, functioned as a T cell modulator and not as a T cell-depleting agent *in vivo* as desired and expected from murine *in vivo* experiments. Similar to low-dose CD28-SA application (14), treatment with a conventional anti-CD28 mAb could be monitored in patients by detecting mAb bound to CD28 and by measuring the overall level of CD28 expression. Treatment of pigs with a CD28-SA differed from infusion of mAb 3D11 in that it led to upregulation of CD28 expression on T cells and induced proliferation of Tconv in peripheral blood (14) (Figure 4). However, infusion of both types of anti-CD28 mAb led to an accumulation of Treg in the infarcted myocardium (14) (Figure 6). Currently, it is unclear whether there are factors in the infarcted myocardium that cause the accumulation of Treg or whether Treg receive a form of ‘licensing’ e.g. in mediastinal lymph nodes after anti-CD28 mAb treatment, influencing their migratory behaviour. Our data add another piece to the understanding of the underlying pathophysiological mechanisms.

The clinically most advanced approach of targeting and blocking CD28 costimulation is the FR104 antibody fragment, which is a pegylated anti-CD28 single chain Fv of the conventional anti-CD28 mAb CD28.3 that blocks ligand binding and that has successfully passed phase II clinical testing (30). Apart from the monovalent FR104, a humanised Fc-silenced anti-CD28 mAb, FK734, has been generated and successfully used in an allogeneic skin graft rejection model in humanised mice (31). For FK734, its capacity to block ligand binding has not been tested or published. Due to the presumed risk associated with bivalent binding of CD28 by intact mAb, further development of FK734 antibody was stopped (32). Our data, however, indicate that binding CD28 bivalently outside of the C’’D loop is not per se associated with adverse effects.

To understand whether treatment of pigs after MI with mAb 3D11 has a positive impact on cardiac function in the mid- and long-term, serial cardiac MRI (33) measurements up to week five after MI would be ideal. Short-term data obtained at day 7 after myocardial infarction do not permit linking alterations in the Treg compartment, predominantly in the border zone, to enhanced cardiac wound healing. However, for now, missing long-term functional outcome data are a key limitation of our study. Another limitation of our study is the low number of pigs in the MOPC-21 mAb-treated control group.

Taken together, our finding that conventional anti-CD28 mAb can be safely administered to pigs, even at a saturating dose, advances the development of drugs targeting T cell costimulation. In particular, it is not yet clear to what degree T cell costimulation should be inhibited in humans to achieve optimal results. Here, the combination of selective inhibition of ligand binding to CD28 and retained agonistic activity, as is the case for mAb 3D11, might strike the right balance between immunomodulation on the one hand and preserved immunosurveillance (26) on the other hand.

## Declarations

## Acknowledgements

The authors would like to thank Franziska Barthelmes, Katja Blouin, Helga Wagner, Nicole Maier, Sandra Werner-Wittig and Jutta Meißner-Weigl for their outstanding and supportive work in the laboratory as well as Verena Burkard and Kerstin Körner for their invaluable help with animal experiments and for their dedicated and loving animal care.

## Data availability

The data supporting the conclusions of this study are provided within the paper and its supplementary information files or can be accessed from the corresponding author upon reasonable request.

## Abbreviations

A4C: apical 4-chamber view
AE: adverse event
BSA: bovine serum albumin
BW: bodyweight
BZ: border zone
CD28-SA: superagonistic monoclonal anti-CD28 antibody clone 4D12
Con A: Concanavalin A
CRS: cytokine release syndrome
DC: dendritic cell
DMSO: dimethyl sulfoxide
DPBS: Dulbecco’s phosphate-buffered saline
DTAC: dodecyltrimethylammonium-chloride
E/A: mitral valve inflow profile
ECG: electrocardiography
E/É: mitral annulus velocities
FACS: fluorescence activated cell sorting
HBSS: Hankś balanced salt solution
IC: infarct core
IL: interleukin
i.m.: intramuscular
i.v.: intravenous
LAD: left anterior descending artery
LVEDV: left ventricular enddiastolic volume
LV-EF: left ventricular ejection fraction
LF-FS: left ventricular fractional shortening
LVIDd: left ventricular internal diastolic diameter
LVOT: left ventricular outflow tract
mAb: monoclonal antibody
MACS: magnetic activated cell sorting
MI: myocardial infarction
M-Mode: motion mode
MOPC-21: isotype control monoclonal antibody
PBMC: peripheral blood mononuclear cells
PBS: phosphate-buffered saline
PFA: paraformaldehyde
PLAX: parasternal long-axis view
RM: remote myocardium
SV: stroke volume
TBS: tris-buffered saline
Tconv: conventional T cells
Treg: regulatory T cells

