## Supplementary Information for "Treatment with saturating dose of conventional anti-CD28 monoclonal antibody well tolerated in pig acute myocardial infarction model"

**Supplemental Figure S1. Coronarangiography and macroscopically visible pathological changes in pig model of myocardial infarction. A**) The LAD and its first lateral branch (blue star) were visualized by coronary angiography. The (upper) image shows the vessel before and (lower) during myocardial infarction induction with an inflated occlusion balloon. **B**) The lighter infarcted area of the pig hearts can be identified macroscopically (colored line) – one example of an explanted heart of a control mAb and of a mAb 3D11-treated animal. **C**) A slice from the left ventricle to illustrate the sampled areas. The three zones (infarct core, border zone and remote myocardium) can be differentiated macroscopically.

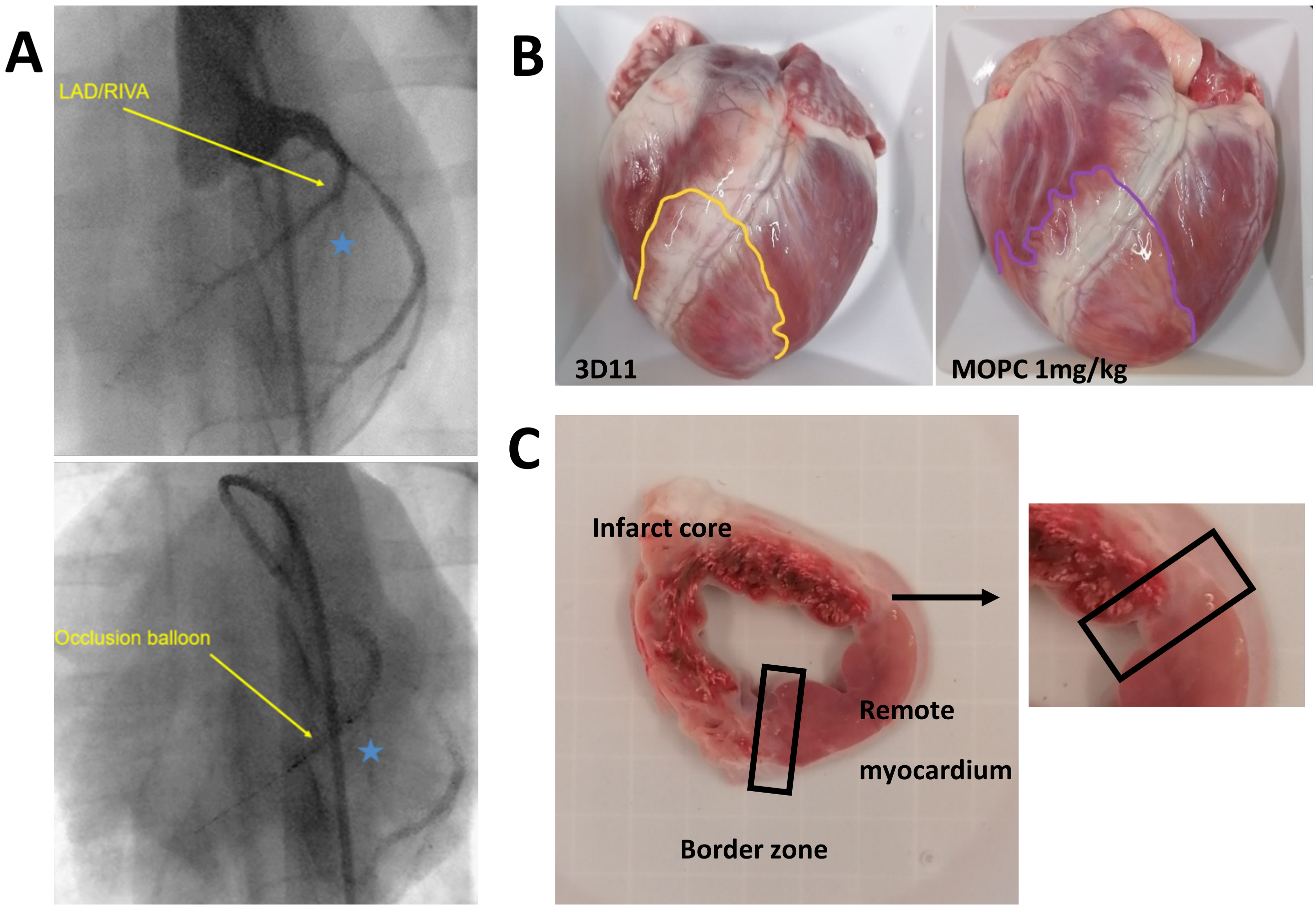

**Supplemental Figure S2.**

**Gating strategy to monitor antibody infusion and T cell subsets in peripheral blood. A**) Identification of CD4$^+$ and CD8$^+$ T cells in peripheral blood. **B**) detection of infused mAb 3D11 on CD4$^+$ T cells on day 7 (black) compared to day 0 (grey). **C**) CD28 expression on peripheral blood CD4$^+$ T cells before (grey) and seven days after MI induction (black). Dashed lines show controls without primary and secondary antibody for detection of CD28 expression. **D**) Detection of TruCount beads (left) and leukocyte gating (right). **E**) Among leukocytes, CD45$^+$ CD3$^+$ T cells were further gated according to CD4 and CD8 expression to determine absolute numbers of CD3$^+$ CD45$^+$ CD4$^+$ and CD8$^+$ T cells per μl of blood. **F**) Detection of CD25$^+$ Foxp3$^+$ Treg among gated CD4$^+$ T cells. **G**) Ki-67 versus Foxp3 expression of CD4$^+$ T cells of peripheral blood.

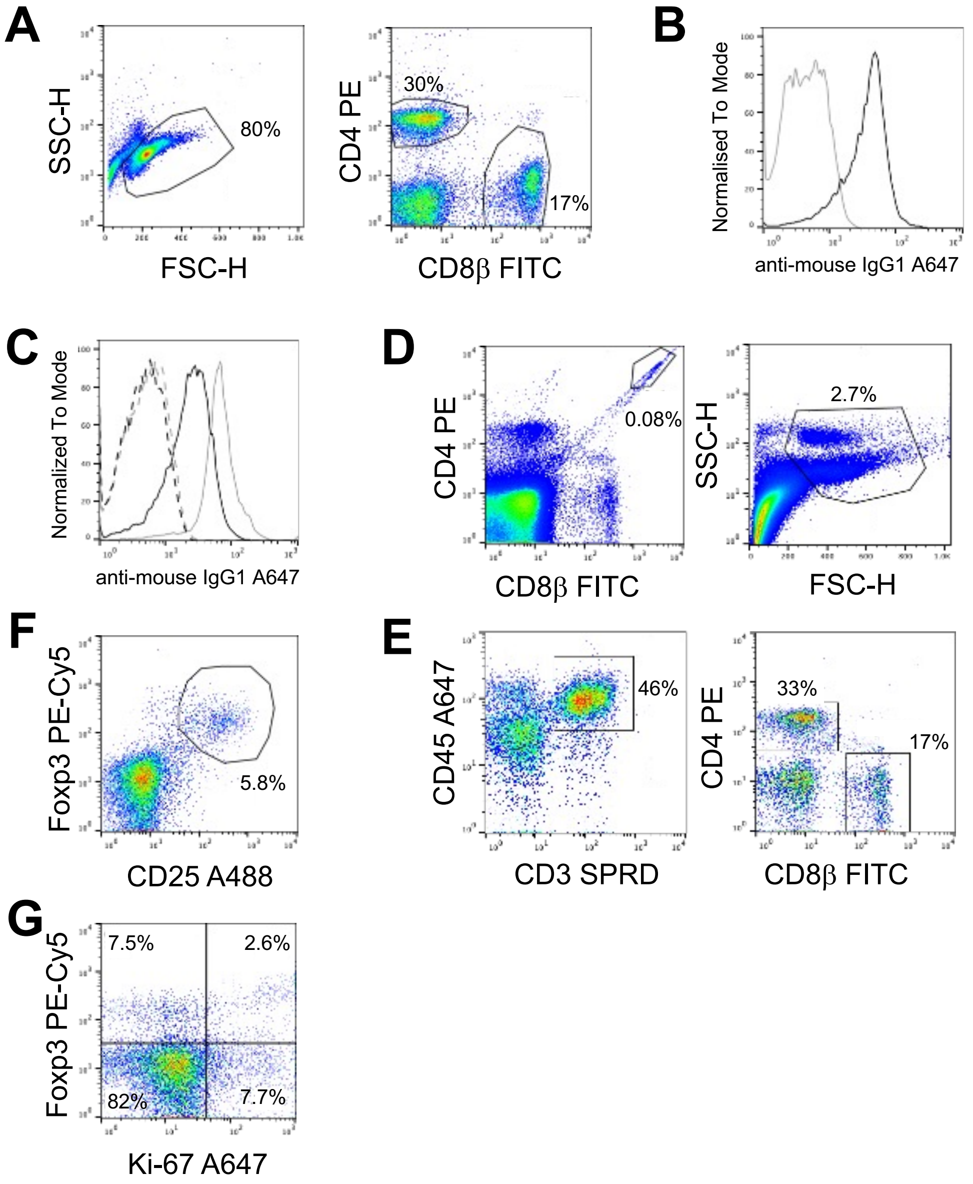

### Supplemental Table 1

**List of monoclonal primary antibodies used for flow cytometry (FC) analysis of whole blood samples**

| **Antigen** | **Conjugate** | **Isotype** | **Clone** | **Species** | **Format** | **Source** | **Concentration** |
|---|---|---|---|---|---|---|---|
| CD3ε | SPRD | IgG1 | PPT3 | mouse:pig | labeled | southern biotech | 1:500 |
| CD4α | PE | IgG2b | 74-12-4 | mouse:pig | labeled | BD | 1:100 |
| CD8β | FITC | IgG1 | PPT23 | mouse:pig | labeled | Biorad | 1:400 |
| CD45 | A647 | IgG1 | K252.1E4 | mouse:pig | labeled | Biorad | 1:15 |

### Supplemental Table 2

**List of monoclonal primary antibodies used for flow cytometry (FC) analysis of PBMC, spleen, lymph node, and heart samples**

| **Antigen** | **Conjugate** | **Isotype** | **Clone** | **Reactivity** | **Format** | **Source** | **Concentration** |
|---|---|---|---|---|---|---|---|
| none | / | all | polyclonal | mouse:unknown | unlabelled | Sigma Aldrich | 1:50 |
| pig CD4α | PE | IgG2b | 74-12-4 | mouse:pig | labelled | BD | 1:100 |
| pig CD8β | FITC | IgG1 | PPT23 | mouse:pig | labelled | Biorad | 1:400 |
| Foxp3 | PE-Cy5 | IgG2a | FJK-16s | rat:mouse | labelled | Thermo Fisher | 1:100 |
| Ki67 | A647 | IgG1 | B56 (RUO) | mouse:human | labelled | BD | 1:300 |
| pig CD25 | A488 | IgG1 | K231.3B2 | mouse:pig | labelled | Biorad | 1:200 |
| mouse IgG | A647 | IgG1 | RM61-1 | rat:mouse | labelled | BioLegend | 1:2000 |
| CD28 | / | $IgG_1$ | 3D11 | mouse:pig | unlabelled | in vivo Biotech | 1:200 |
| CD28 | / | $IgG_1$ | 4D12 | mouse:pig | unlabelled | in vivo Biotech | 1:100 |

### Supplemental Table 3

#### Flow cytometry staining workflow

| Row / Panel | 20 min on ice | Wash 1 | 15 min on ice | Wash 2 | 15 min on ice | 15 min on ice | Wash 3 | 30 min on ice | Wash 4 | 45 min at RT (in the dark) | Wash 5 |
|---|---|---|---|---|---|---|---|---|---|---|---|
| **1** | --- | 1x, change well; 1x | amIg-A647 | 1x | normal mouse Ig (nmIgG) | CD8β-FITC CD4-PE | 2x | | | | |
| **2** | 4D12 | 1x, change well; 1x | amIg-A647 | 1x | normal mouse Ig (nmIgG) | CD8β-FITC CD4-PE | 2x | | | | |
| **3** | 3D11 | 1x, change well; 1x | amIg-A647 | 1x | normal mouse Ig (nmIgG) | CD8β-FITC CD4-PE | 2x | | | | |
| **4** | --- | | --- | 1x | normal mouse Ig (nmIgG) | CD8β-FITC CD4-PE CD27-APC | 2x | | | | |
| **5** | | | | 1x | normal mouse Ig (nmIgG) | CD25-A488 CD4-PE | 1x PBS | Fix/Perm | 1x | FoxP3-PE-Cy5 Ki67-A647 | 2x |
| **6** | | | | 1x | normal mouse Ig (nmIgG) | CD4-PE | 1x PBS | Fix/Perm | 1x | FoxP3-PE-Cy5 Ki67-A647 | 2x |
| **7** | | | | 1x | normal mouse Ig (nmIgG) | CD4-PE | 1x PBS | Fix/Perm | 1x | --- | 2x |
| **Volumes** | 50 µl | 150 µl FACS-buffer | 50 µl | 150 µl FACS-buffer | 25 µl | 25 µl | 150 µl | 100 µl | 100 µl Perm-buffer | 50 µl | 150 µl FACS-buffer |
